# Identification of pre-synaptic density networks using SV2A PET imaging and ICA in healthy and diseased mice

**DOI:** 10.1101/2022.10.19.512761

**Authors:** Jordy Akkermans, Franziska Zajicek, Alan Miranda, Mohit Adhikari, Daniele Bertoglio

**Affiliations:** Molecular Imaging Center Antwerp (MICA), University of Antwerp, Belgium; Bio-Imaging Lab, University of Antwerp, Belgium

**Author notes:** **Corresponding author:** Prof. Daniele Bertoglio, PhD, University of Antwerp, Universiteitsplein 1, Wilrijk, Belgium.

**Keywords:** Synaptic vesicle glycoprotein 2A, [^11^C]UCB-J, positron emission tomography, rodent, independent component analysis, Huntington’s disease

## Abstract

**Background:** Synaptic vesicle glycoprotein 2A (SV2A) is a vesicle glycoprotein involved in neurotransmitter release. SV2A is located on the pre-synaptic terminals of neurons and visualized using the radioligand [^11^C]UCB-J and positron emission tomography (PET) imaging. Thus, SV2A PET imaging can provide a proxy for pre-synaptic density in health and disease. This study aims to apply independent component analysis (ICA) to SV2A PET data acquired in mice to identify pre-synaptic density networks (pSDNs), explore how ageing affects these pSDNs, and determine the impact of a neurological disorder on these networks.

**Methods:** We used [^11^C]UCB-J PET imaging data (*n*=135) available at different ages (3, 7, 10, and 16 months) in wild-type (WT) C57BL/6J mice and in diseased mice (mouse model of Huntington’s disease, HD). First, ICA was performed on a healthy dataset after it was split into two equal-sized samples (*n*=36 each) and the analysis was repeated 50 times in different partitions. We tested different model orders (8, 12, and 16) and identified the pSDNs. Next, we investigated the effect of age on the loading weights of the identified pSDNs. Additionally, the identified pSDNs were compared to those of diseased mice to assess the impact of disease on each pSDNs.

**Results:** Model order 12 resulted in the preferred choice to provide six reliable and reproducible independent components (ICs) as supported by the cluster-quality index (I_Q_) and regression coefficients (β) values. Temporal analysis showed age-related statistically significant changes on the loading weights in four ICs. ICA in an HD model revealed a statistically significant disease-related effect on the loading weights in several pSDNs in line with the progression of the disease.

**Conclusion:** This study validated the use of ICA on SV2A PET data acquired with [^11^C]UCB-J for the identification of cerebral pre-synaptic density networks in mice in a rigorous and reproducible manner. Furthermore, we showed that different pSDNs change with age and are affected in a disease condition. These findings highlight the potential value of ICA in understanding pre-synaptic density networks in the mouse brain.

## 1. Introduction

Synaptic vesicle glycoprotein 2A (SV2A) is the most widely distributed transmembrane glycoprotein present on secretory vesicles in the pre-synaptic terminal of neurons throughout the central nervous system (Bajjalieh et al., 1994). SV2A is involved in the regulation of neurotransmission (Feany et al., 1992) and is a target for the antiepileptic drug levetiracetam (Lynch et al., 2004). SV2A can be used as a marker to visualize pre-synaptic density distribution *in vivo* using positron emission tomography (PET) imaging thanks to the SV2A radioligands available, including [^11^C]UCB-J (Nabulsi et al., 2016). [^11^C]UCB-J displayed optimal imaging characteristics to quantify and monitor synaptic density (Finnema et al., 2016; Finnema et al., 2018) and can be used to visualize SV2A loss in neurological disorders including Alzheimer’s Disease (AD), Parkinson’s Disease (PD), and Huntington’s Disease (HD), as well as other neurodegenerative and neuropsychiatric disorders (Chen et al., 2018; Delva et al., 2020; Delva et al., 2022; Matuskey et al., 2020).

In addition to human application, [^11^C]UCB-J displayed excellent imaging characteristics in rodents and non-human primates (Bertoglio et al., 2020; Nabulsi et al., 2016; Thomsen et al., 2021). Thus, [^11^C]UCB-J PET imaging can be applied in a preclinical setting to model neurological and neuropsychiatric disorders as demonstrated by recent studies by our and other groups (Bertoglio et al., 2022a; Bertoglio et al., 2022b; Glorie et al., 2020; Toyonaga et al., 2019)

Given the brain-wide distribution of SV2A, regional analysis of SV2A PET data may be limiting the amount of information that can be obtained. In this context, data-driven approaches such as independent component analysis (ICA), a blind source separation technique, can separate the brain signal into distinct components, here defined as pre-synaptic density networks (pSDNs), without having any knowledge beforehand about the source signals (Bell and Sejnowski, 1995). ICA divides the data into different independent components (ICs) which are assumed to be statistically independent with a non-gaussian distribution (Hyvärinen and Oja, 2000). ICA is most applied to functional magnetic resonance imaging (fMRI) data to identify networks that show correlated neuronal activity (McKeown et al., 1998), although few studies applied the technique to PET imaging as well. Recently, Fang and colleagues (Fang et al., 2021) applied for the first time ICA to [^11^C]UCB-J PET data in healthy humans, demonstrating the feasibility of the approach.

Starting from this evidence, we aimed at expanding the approach to the preclinical domain by applying ICA to [^11^C]UCB-J PET in mice. First, the technique was validated by generating robust pSDNs and identifying their characteristics in mice. Next, we explored whether the pSDNs would be affected with ageing at 3, 7, 10, and 16 months. Lastly, we studied the potential value of pSDNs as marker of disease condition in a mouse model of HD, a neurodegenerative disorder known to affect SV2A levels (Bertoglio et al., 2022b; Delva et al., 2022).

## 2. Material and Methods

### 2.1. Animals

Male wild-type (WT) C57BL/6J mice and diseased zQ175DN (mouse model of HD) littermates with a C57BL/6J background were included (Heikkinen et al., 2012; Menalled et al., 2012). Two experimental groups were used: a cross-sectional group at 3 months (*n*=16 per condition) and a longitudinal group imaged at 7 (*n*=17, WT; *n*=18, HD), 10 (*n*=23, WT; *n*=15, HD), and 16 (*n*=16, WT; *n*=14, HD) months (**Supplementary Figure 1**). Data analysis for the current study was based on previously acquired data (Bertoglio et al., 2022b). All animals were purchased from Jackson Laboratories (Bar Harbour, Maine, USA). Before the start of the experiments, the animals had at least one week to acclimatize single-housed in individually ventilated cages under a 12-hour light/dark cycle. The environment was temperature- and humidity-controlled, food and water were given *ad libitum*. All procedures were performed following the European Committee Guidelines (decree 2010/63/CEE) and were approved by the Ethical Committee for Animal Testing (ECD 2017-27) at the University of Antwerp.

**Figure 1:**
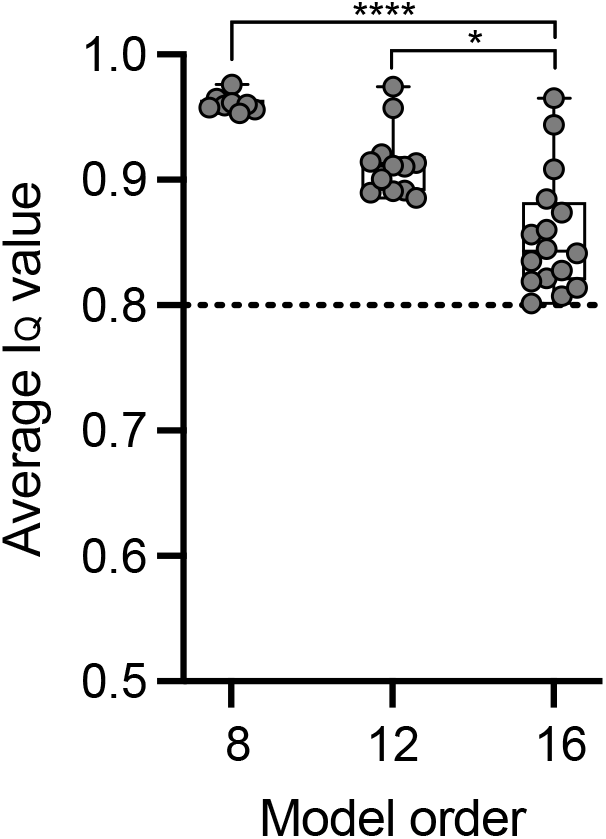
Cluster-quality index (I_Q_) for different model orders. The dashed line represents the threshold of 0.8 for reliable ICs. Box = median and quartiles, whiskers = min to max values. \**p*<0.05, \*\*\*\**p*<0.0001.

### 2.2. Tracer synthesis

[^11^C]UCB-J synthesis was accomplished on an automated synthesis module (Carbonsynthon I, Comecer, The Netherlands) in the Antwerp University Hospital, adapting the previously reported process (Nabulsi et al., 2016) to the system (Bertoglio et al., 2020). As previously published the average radiochemical purity was greater than 99%, and molar activity was 96.5 ± 13.3 GBq/μmol (Bertoglio et al., 2022b).

### 2.3. PET procedures

The PET scans were conducted using two Siemens Inveon PET-CT scanners (Siemens Preclinical Solution, USA). Before each scan, the animals were weighed and anaesthetized with inhalation of 5% Isoflurane in oxygen. A catheter was used to intravenously administer the tracer into the tail vein. 1.5% Isoflurane was used for anaesthesia maintenance. During the scan, parameters such as respiration and temperature were continuously monitored using a Monitoring Acquisition Module (Minerve, France). An intravenous bolus injection (5.4 ± 1.3 MBq) of [^11^C]UCB-J was administered over 12 seconds using an automated pump (Pump 11 Elite, Harvard Apparatus, USA) at the start of the scanning procedure. Following the PET scan, a CT scan (10 min 80 kV/500 μA) was performed for attenuation correction.

### 2.4. Image processing and quantification

PET data is collected and histogrammed before being reconstructed into 33 frames of increasing length (12×10s, 3×20s, 3×30s, 3×60s, 3×150s, and 9×300s). All images were reconstructed in 8 iterations and 16 subsets using the 3D ordered subset expectation maximization (OSEM 3D) algorithm utilizing a list-mode iterative reconstruction with proprietary spatially variant resolution modeling for quantitative analysis (Miranda et al., 2020). Corrections for normalization, dead time, and CT-based attenuation were applied. PET image frames were reconstructed using 0.776×0.776×0.796 mm^3^ voxels on a 128×128×159 grid.

PMOD 3.6 software (Pmod Technologies, Zurich, Switzerland) was used to analyze and process the PET data. Kinetic modeling was performed as previously described (Bertoglio et al., 2022b). Specifically, parametric *V*_T (IDIF)_ maps needed for the ICA were generated through voxel-wise Logan graphical analysis, whereby the input function was accounted for by using a non-invasive IDIF to acquire *V*_T (IDIF)_.

### 2.5. Independent component analysis

ICA was performed in the source-based morphometry (SBM) module of the GroupICA toolbox GIFT. The simplest version of the model can be described as:

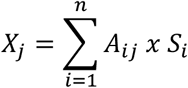

With *X* representing the input PET data matrix, *A* the mixing matrix, and *S* the source matrix, with *A* and *S* being the output of the analysis. The input matrix (*X*) contains information about the voxels for each subject (*j*). The mixing matrix gives information about the loading weights for each subject for each identified component (*i*). The loading weights of the subjects are used as a measure of contribution for a given IC, they are used to determine the differences between genotypes. The source matrix contains information about the voxel distribution on a group-level for each IC (*i*), which form the source maps (*S*) (Bell and Sejnowski, 1995; Hyvärinen and Oja, 2000). Before starting ICA, a centering step was performed as part of the principal component analysis (PCA). The centering of the data was done by calculating the mean *V*_T (IDIF)_ value inside a mask of the whole brain with Matlab (2015a). Next, demeaning of the *V*_T (IDIF)_ maps was performed, i.e. the mean *V*_T (IDIF)_ of the brain inside the mask was subtracted for each subject, to remove the effect of subject variability.

In order to identify robust ICs, the reliability and repeatability were evaluated at three different model orders (number of ICs),namely 8, 12, and 16. The setup of the analysis was matched to the settings used by Fang and colleagues (Fang et al., 2021). The analysis was done in healthy mice using two separate samples (*n*=36 each) for any given model order. Initially, two masks were used to delineate a region of interest (ROI), a mask of the whole brain and a mask excluding the cerebellum. The latter mask was included to evaluate if other components would perform differently since the cerebellum was our most reliable IC and it is found to be anti-correlated to cortical regions as measured with [^18^F]FDG-PET and rs-fMRI (Tomasi et al., 2017). Since the impact of the cerebellar IC on other components was negligible, we reported the results obtained using only the whole brain mask. The results were scaled to z-score and the InfoMax algorithm was used (Bell and Sejnowski, 1995).

The repeatability of the ICs was evaluated with the ICASSO module. This module repeats the extraction of the ICs from a single run 50 times to measure the repeatability of the components. The repeatability was represented as the cluster-quality index (I_Q_). ICs with an I_Q_ value above 0.8 are considered repeatable (Fang et al., 2021). To assess the reliability, the ICs were compared between samples within each run. Based on the spatial similarities with multiple regression, these similarities were expressed as beta values. The pair with the highest β-value was selected and visually compared until the ICs did not match anymore. Across model orders, a pattern was identified where everything below a β-value of 0.3 no longer consistently matched. As a result, only ICs exceeding the threshold of β = 0.3 were included. This setup was repeated 50 times with the same settings but in different equal-sized groups to make sure the identification is not specific to a set partition of the dataset.

The components were sorted based on IC spatial similarity across the 50 runs for each model order to make results more easily interpretable. In Matlab, the run with the highest β-values across the components was chosen as reference run. The other runs were reorganized to match the order of ICs of the reference run. The β and I_Q_ values were averaged across the runs and the reliable and repeatable ICs were chosen based on the components whose average reliability & repeatability indices passed the threshold of β = 0.3 and I_Q_ = 0.8. In addition, the β and I_Q_ values of the ICs were compared between model orders to determine the most optimal model order for the exploration of the pSDNs. A summary of the step-by-step procedure is presented in (**Supplementary Figure 2)**. The identified ICs were categorized based on the localization of the signal over the anatomical MRI study template (Bertoglio et al., 2022b) in PMOD. The identification of the components was performed using the Paxinos mouse brain atlas (Paxinos and Franklin, 2003).

Next, in order to investigate whether ageing would affect the ICs, the entire WT dataset was used (*n*=72) and ICA was repeated, in one single run, at the model order 12. The loading weights were grouped per age and extracted for each IC.

Finally, to explore whether pSDNs are affected in a disease condition, the entire dataset of WT and HD mice (*n*=135) was used. The analysis was run in 50 different randomized split samples, to determine the ICs with the new sample size. The loading weights of these ICs were extracted and compared between genotypes to determine age and genotypic differences.

### 2.6. Statistical analysis

The I_Q_ values and loading weights were tested for normality with the Shapiro-Wilk test. The Kruskal-Wallis test was performed to assess the difference between mean loading weights across four age groups for each component as well as on I_Q_ values. Two-way ANOVA was used to evaluate the effect of ageing and disease on the loading weights from both groups. The results were corrected for multiple comparisons with the Bonferroni method. Outliers were excluded if they exceeded 1.96 times the standard deviation (SD) of the group. The statistical analyses were performed with GraphPad Prism (v9.1) statistical software. Data are reported as mean ± SD, unless stated otherwise. All tests were two-tailed and statistical significance was set at *p*<0.05.

## 3. Results

### 3.1. Model order selection

With the goal of determining the most suited model order, repeatability and reliability of the identified components were evaluated and compared between model orders 8, 12, and 16. The repeatability of the components for each model order was evaluated based on the I_Q_ values calculated per component and averaged across the 50 runs. The I_Q_ values for all ICs passed the threshold of 0.8 across all model orders, however, an increase in model order showed a statistically significant decrease in I_Q_ value (model order 8: 0.96 ± 0.05, model order 12: 0.93 ± 0.07, model order 16: 0.91 ± 0.09; *p*<0.0001) (**Figure 1** and **Table 1**).

**Table 1:** β and I_Q_ values for each IC identified in healthy mice at model order 8, 12 and 16. S1BF = Primary somatosensory barrel field. Values are presented as mean ± SD. ^#^These components did not pass the 0.3 threshold, so they were not considered further in the analysis.

| IC | Pre-synaptic density network | Model order 8 |  | Model order 12 |  | Model order 16 |  |
| --- | --- | --- | --- | --- | --- | --- | --- |
| | | $\beta$ | $I_Q$ | $\beta$ | $I_Q$ | $\beta$ | $I_Q$ |
| IC1 | Cerebellar | $0.78 \pm 0.06$ | $0.98 \pm 0.04$ | $0.79 \pm 0.03$ | $0.97 \pm 0.02$ | $0.75 \pm 0.04$ | $0.97 \pm 0.05$ |
| IC2 | Medulla & pons | $0.63 \pm 0.24$ | $0.96 \pm 0.05$ | $0.69 \pm 0.13$ | $0.96 \pm 0.05$ | $0.69 \pm 0.10$ | $0.94 \pm 0.05$ |
| IC3 | Septal nucleus | $0.39 \pm 0.24$ | $0.96 \pm 0.06$ | $0.47 \pm 0.16$ | $0.92 \pm 0.09$ | $0.47 \pm 0.12$ | $0.91 \pm 0.10$ |
| IC4 | Temporal lobe | $0.37 \pm 0.31$ | $0.96 \pm 0.06$ | $0.43 \pm 0.27$ | $0.91 \pm 0.08$ | $0.20 \pm 0.16^{\#}$ | $0.81 \pm 0.13$ |
| IC5 | Left striatal | $0.32 \pm 0.26$ | $0.96 \pm 0.04$ | $0.33 \pm 0.15$ | $0.91 \pm 0.10$ | $0.33 \pm 0.12$ | $0.86 \pm 0.12$ |
| IC6 | S1BF | $0.32 \pm 0.16$ | $0.95 \pm 0.07$ | $0.21 \pm 0.19^{\#}$ | $0.89 \pm 0.10$ | $0.25 \pm 0.14^{\#}$ | $0.85 \pm 0.13$ |
| IC7 | Upper cortex | $0.10 \pm 0.28^{\#}$ | $0.96 \pm 0.05$ | $0.33 \pm 0.22$ | $0.91 \pm 0.10$ | $0.41 \pm 0.14$ | $0.88 \pm 0.13$ |

The reliability of the ICs was assessed by averaging the β values calculated with multiple regression across the 50 runs for each model order. Both model order 8 and 12 could identify 6 ICs reliably, with the latter having slightly better β-values and lower standard deviations (β=0.47 ± 0.22 at model order 8 and β=0.50 ± 0.16 at model order 12), whereas model order 16 could reliably classify only 5 ICs. Based on fewer robust ICs in model order 16 and the decreasing I_Q_ values, there was no indication of a gained benefit when using a higher model order. Thus, model order 12 was chosen based on the reliability and repeatability of the ICs.

### 3.2. Identification of pre-synaptic density networks in mice

Classification of the identified IC was done based on the reliability measurements initially detected in model order 8. Component were labelled based on the anatomical structures covered by the peak of IC. The most reliable IC identified at model order 12 was the cerebellar component (IC1), which was followed by the IC2, mainly located in the pons and medulla area. IC3 was concentrated around the septal nucleus area, whereas IC4 covered the temporal lobe bilaterally. The fifth component identified the caudal left striatum. IC6 was visualized as cortical component, however it did not pass the 0.3 β threshold when using model order 12. Finally, IC7 extended across the entire top cortical structures and was labelled as upper cortex. An overview of the identified pSDNs using model order 12 is shown in **Figure 2**.

**Figure 2:**
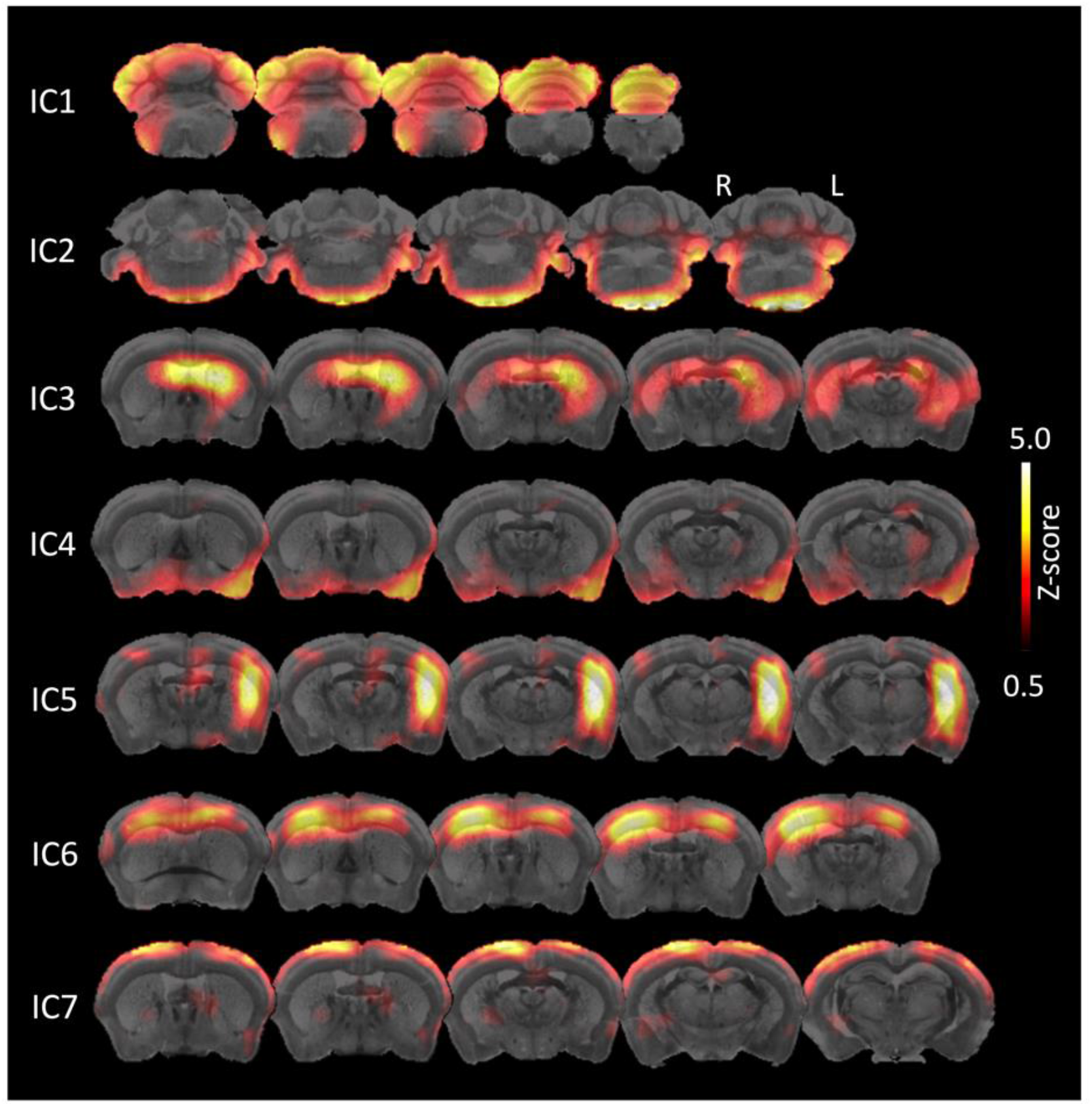
Visualization of the pSDNs identified in healthy mice (*n*=72) using model order 12. Different coronal slices overlaid onto the MRI template are presented for each component. The scale is shown on the right with a threshold of Z ≥ 0.5. The components were extracted from a separate analysis done with the full dataset (*n*=72) for visualization.

When comparing across model orders, model order 8 and 12 both identified 6 reliable (β > 0.3) ICs, five of which matched (IC1-IC5). The primary somatosensory barrel field (S1BF) (IC6) (β = 0.32 ± 0.16) was reproduced in model order 8 but was not reliably estimated in model order 12 (β = 0.21 ± 0.19). Similarly, the upper cortex (IC7) could not be reproduced in model order 8 (β = 0.10 ± 0.28) but was considered reliable in model order 12 (β = 0.33 ± 0.22). When increasing to model order 16, only five ICs were identified robustly, all of them found in the previous model orders (IC1-IC3, IC5, IC7). The temporal lobe (IC4) (β = 0.20 ± 0.16) from model order 12 and S1BF (IC6) (β = 0.25 ± 0.14) from model order 8 were not reproduced. Across model orders the cerebellar was the most reliably identified (β = 0.78 ± 0.06; 0.79 ± 0.03; and 0.75 ± 0.04 at model order 8, 12, and 16, respectively). An overview of the robustly identified ICs are presented in **Table 1**, whereas for visual comparison see **Supplementary Figure 3 and 4**.

### 3.3. Evolution of pre-synaptic density networks with age

To determine whether ageing influences pSDNs, the entire dataset of healthy mice (*n*=72) was used to extract the loading weights at model order 12. A significant ageing effect was observed for four out of the six ICs (**Figure 3**). Three of those ICs had a positive age effect: cerebellar (IC1, *K-W statistic* = 12.03, *p*=0.0073), medulla & pons (IC2, *K-W statistic* = 35.07, *p*<0.0001), temporal lobe (IC4, *K-W statistic* = 32.19, *p*<0.0001). On the other hand, the upper cortex (IC7) was the only component showing a negative age effect, *K-W statistic* = 27.70, *p*<0.0001. The septal nucleus (IC3) and left striatal (IC5) did not show a significant age effect (*K-W statistic* = 4.538, *p*=0.2089 and *K-W statistic* = 4.529, *p*=0.2097, respectively).

**Figure 1:**
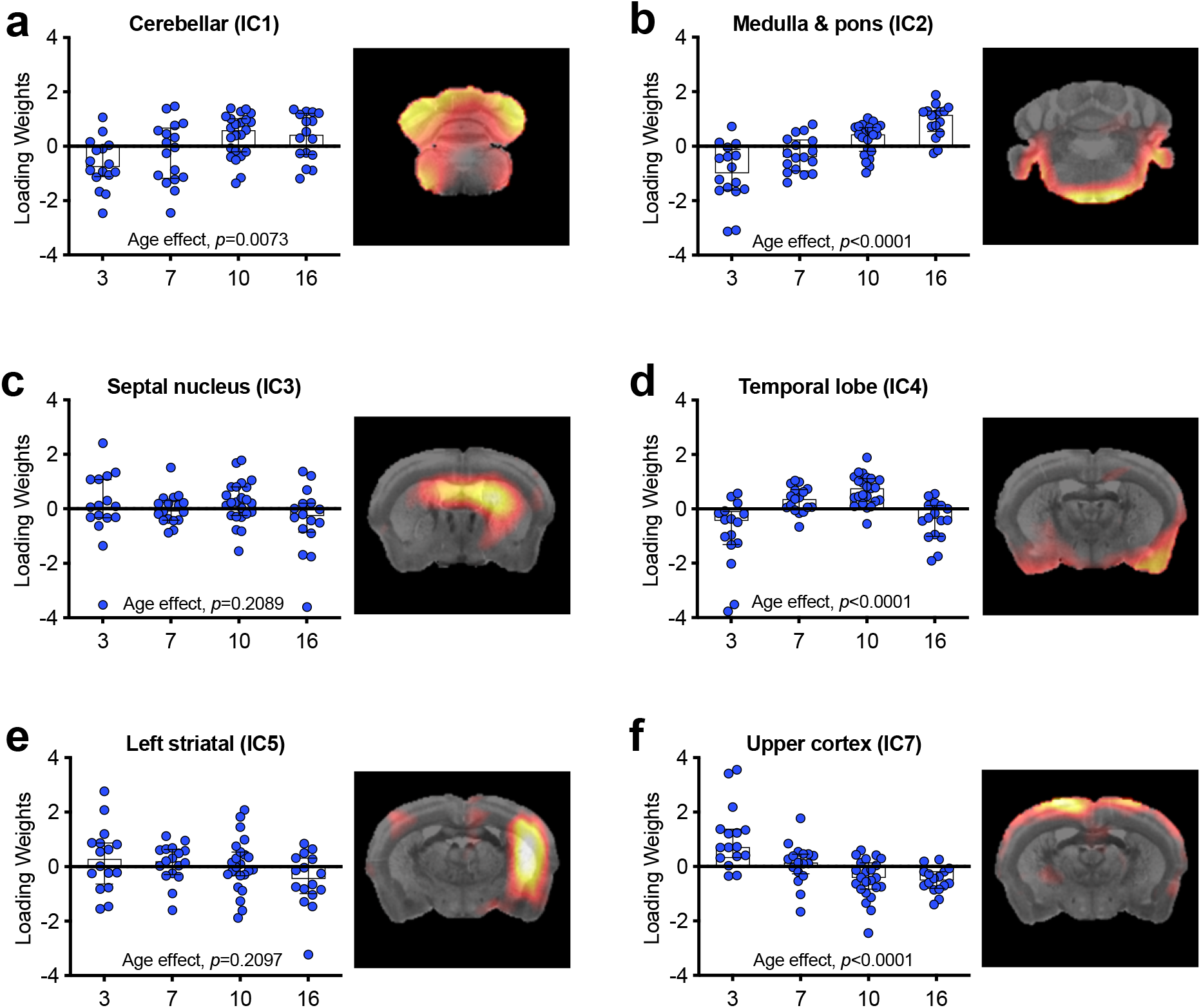
Age effect for every IC identified in healthy mice at model order 12. Each graph shows the age-effect, based on the loading weights, across the four time points. The graphs are accompanied by a visual representation of the ICs in coronal plane. Data are shown median ± interquartile ranges.

### 3.4. Disease condition affects pre-synaptic density networks in mice

The effect of a disease condition, HD, on the pSDNs was assessed by pooling the dataset of healthy and HD mice together (*n*=135) using model order 12. On one hand, the increased sample size resulted in the identification of four additional ICs (IC6, S1BF; IC8, olfactory bulb; IC9, midbrain, and IC10, right rhinal cortex) passing the β-value threshold. On the other hand, the left striatal component (IC5) which was previously identified at model order 12 was not detected anymore. An overview of the ICs is presented in **Supplementary Table 1** and **Supplementary Figure 5**. Statistical analysis confirmed the significant age effect in most ICs, with the exception of the septal nucleus (IC3, *F*_(3,127)_ = 0.5376, *p*=0.6574) and the S1BF (IC6, *F*_(3,127)_ = 0.2862, *p*=0.8353) components (**Supplementary Table 2**).

Comparison of the loading weights between healthy and HD mice identified a significant disease effect for all ICs with the exclusion of the septal nucleus (IC3, *F*_(1,127)_ = 1.115, *p*=0.2931) and the S1BF (IC6, *F*_(1,127)_ = 3.555, *p*=0.0617) components (**Figure 4** and **Supplementary Table 2**). After multiple comparisons correction for pair-wise comparisons, none of the pSDNs was affected by disease condition at 3 months of age. Conversely, as of 7 months of age till 16 months of age, a significant difference in pSDNs loading weights between healthy and HD mice was found in seven out of eight ICs (**Figure 4** and **Supplementary Table 2**).

**Figure 2.**
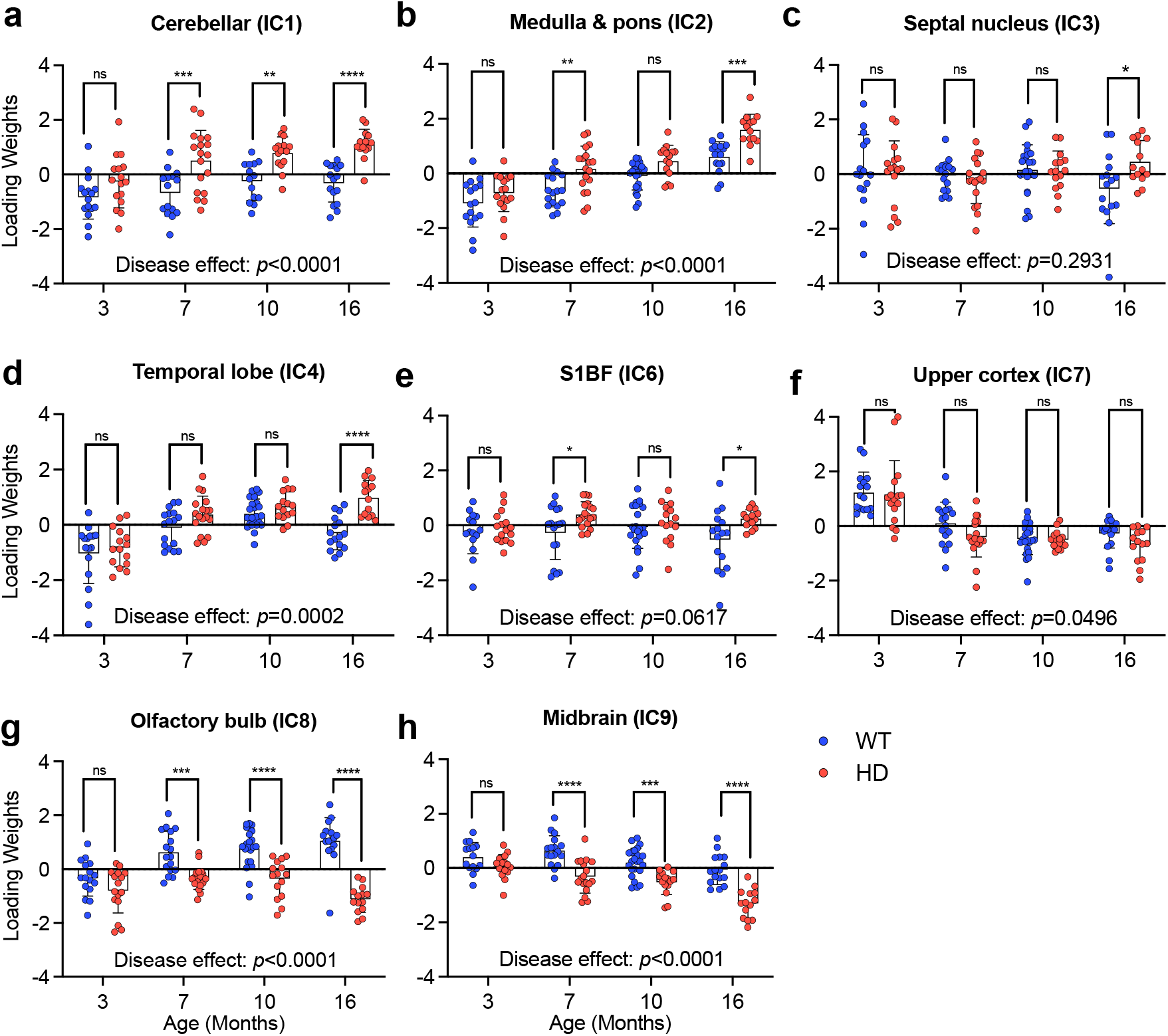
Comparison of loading weights for the ICs identified between healthy and diseased mice (*n*=135). S1BF = Primary somatosensory barrel field (IC6). The right rhinal cortex (IC10) is not analyzed since it did not appear in the separate analysis. ^ns^*p*>0.05, \**p*<0.05, \*\**p*<0.01, \*\*\**p*<0.001, \*\*\*\**p*<0.0001.

## 4. Discussion

In this study, we validated the use of ICA on SV2A PET data to identify cerebral pre-synaptic density networks in mice in a rigorous and reproducible manner. First, we tested three different model orders (namely 8, 12, and 16) and identified the corresponding ICs. Next, we showed that the loading weights of the identified pSDNs are affected by ageing and that a disease condition, such as HD, has an impact on the loading weights of several pSDNs.

The methodology used during the study was based on the approach used in humans by Fang and colleagues (Fang et al., 2021). In addition to the established method, we repeated the runs 50 times in different partitions of the dataset with the objective of ensuring results were not tied to a specific division and that the components could be robustly identified.

Initially, we compared different model orders (8, 12, and 16) to select the optimal number of source networks. A total of six ICs were identified using model orders 8 and 12, while increasing to model order 16, led to fewer robust ICs and decreasing I_Q_ values. In addition, model order 8 displayed less stable β-values, thus model order 12 was selected as optimal choice. Despite the constrain to β-values above 0.3, we could still repeatedly detect the same ICs to be extracted at different model orders. A similar observation was previously reported in humans (Fang et al., 2021). However, whereas the observed six ICs shared similarities with the human ones reported by Fang and colleagues (Fang et al., 2021), in their work they were able to identify 13 ICs at model order 18. It is not surprising that a higher model order and more ICs could be detected in humans given the better spatial resolution in relation to the brain. Typically, a preclinical small-animal PET scan has around three times better resolution than a clinical PET scan, however, due to the difference in brain size, the preclinical PET scan would have to be 15 times better to be on the same level (Kuntner and Stout, 2014; Yao et al., 2012).

ICA is ‘classically’ applied in fMRI data based on blood oxygenation-level dependent (BOLD) contrast for the identification of resting state networks (RSNs), thus a methodological comparison of the outcome compared to SV2A-based pSDNs is intuitive. The identified pSDNs show some similarity in locations compared to RSNs in mice (Zerbi et al., 2015). For instance, the septal nucleus (IC3), left striatal (IC5), S1BF (IC6) and the olfactory bulb (IC8) pSDNs are similarly found in RSNs, with the striatal component being bilateral in the RSNs. Other components, i.e. the cerebellar (IC1), and the upper and right rhinal cortices (IC7, IC10) were split into multiple networks in the RSNs (Zerbi et al., 2015). Despite certain similarities, differences are to be expected given the intrinsic methodological difference between the approaches. Although both fMRI and SV2A PET are measuring neuronal activity, they focused on profoundly different aspects. Using the BOLD contrast, fMRI highlights changes in vascular activity directly or indirectly related to neuronal activity. In contrast, [^11^C]UCB-J PET quantifies SV2A protein at the pre-synaptic cleft, thus provides a more structural measurement of synaptic density. The structural relevance of SV2A PET is further supported by a recent fMRI and [^11^C]UCB-J PET study with visual-activation tasks (Smart et al., 2021) to determine the effect of neural activity on [^11^C]UCB-J binding. Authors reported the *V*_T_ and binding potential of the tracer remained unchanged during the task, whereas a change in tissue influx *K*_1_ in the primary visual cortex correlating with the BOLD fMRI response following visual stimuli was measured (Smart et al., 2021). As such, future studies applying ICA to [^11^C]UCB-J parametric *K*_1_ maps in comparison to BOLD fMRI are warranted to understand whether [^11^C]UCB-J might entail a both structural and functional network information.

Based on the regional analysis of [^11^C]UCB-J *V*_T (IDIF)_ data, we reported a significant age effect (Bertoglio et al., 2022b), thus, one of the objectives of this study was to investigate whether a similar effect applies to the identified pSDNs. Starting from the six identified pSDNs, four of them displayed a significant age effect. Intriguingly, two ICs showed a positive change with age (cerebellar, IC1 and medulla & pons, IC2), while the temporal lobe (IC4) displayed a biphasic and the upper cortex component (IC7) a negative profile.

The temporal changes in the loading weights of these ICs might be associated with altered SV2A density, which was previously shown indirectly in the regional analysis of SV2A PET data (Bertoglio et al., 2022b). Such divergent profiles were also reported in humans (Fang et al., 2021) and may be a consequence of the changes in local variance occurring during ageing as postulated by Fang and colleagues (Fang et al., 2021).

Another aim of this work was to explore the effect of a disease condition (HD) affecting SV2A to the identified pSDNs. In our previous region-based investigation, we demonstrated SV2A based on [^11^C]UCB-J PET data to be significantly reduced in striatum, motor cortex, hippocampus,and thalamus as of 7 months of age with a progressive loss with disease (Bertoglio et al., 2022b). Similarly, analysis of the loading weights in diseased animals demonstrated significant differences in several pSDNs as of 7 months of age with a progressive change, which, intriguingly, displayed both positive and negative changes compared to healthy animals. This represents initial evidence of a disease-related effect on SV2A pSDNs. Although no other work has been published yet, a recent conference abstract suggests effort towards application of SV2A ICA in disease conditions is being made for human studies as well. In this report, authors indicated changes in different ICs for the AD population (Fang et al., 2020). This is consistent with the previous regional observation that SV2A density is reduced in AD patients (Chen et al., 2018).

Collectively, our preclinical work as well as clinical findings are providing initial encouraging evidence of the applicability and utility of pSDNs in neurological and neuropsychiatric diseases. In the near future, it will be interesting to expand this approach to cross-species (e.g. people with HD given the reported loss of SV2A (Delva et al., 2022), as well as to other disorders known to affect SV2A, both preclinically (Bertoglio et al., 2021; Bertoglio et al., 2022a; Glorie et al., 2020) and clinically (Chen et al., 2018; Finnema et al., 2020; Holmes et al., 2019; Matuskey et al., 2020).

Although [^11^C]UCB-J is the most widely used and validated radioligand to target SV2A, it is limited by the short half-life of C11 (20.4 minutes) as well as the technical demanding radiosynthesis, which restricts broad distribution and application. Several F18 bearing radioligands, including [^18^F]SynVesT-1 (Bertoglio et al., 2022c; Naganawa et al., 2021), have been described in the recent years given the more favorable half-life (109.7 minutes), high positron decay and lower positron range. Due to its properties, [^18^F]SynVesT-1 may represent an interesting alternative to [^11^C]UCB-J for the identification and investigation of pSDNs. Future work comparing the outcome of the ICA in the same subject will be of relevance to understand whether either of the radioligands provides better performance.

## 5. Conclusion

This study validated the use of ICA on SV2A PET data acquired with [^11^C]UCB-J for the identification of cerebral pre-synaptic density networks in mice in a rigorous and reproducible manner. This work showed different pSDNs change with age and are affected by disease condition. These findings highlight the potential value of ICA in understanding pre-synaptic density networks in the mouse brain.

## Data availability

Data are available upon request to the corresponding author.

## Supplementary Materials

Supplementary material associated with this article can be found in the online version.

## CRediT Author Contribution Statement

**Jordy Akkermans:** Formal analysis; Methodology; Validation; Visualization; Writing - original draft; Writing - review & editing. **Franziska Zajicek:** Methodology; Writing - review & editing. **Alan Miranda:** Software; Writing - review & editing. **Mohit Adhikari:** Conceptualization; Methodology; Validation; Software; Visualization; Writing - review & editing. **Daniele Bertoglio:** Conceptualization; Funding acquisition; Formal analysis; Methodology; Validation; Visualization; Writing - original draft; Writing - review & editing.

